# Modular inhibitory coding in binary networks

**DOI:** 10.1101/2025.09.22.677833

**Authors:** Bofang Wang, Michal Zochowski

## Abstract

We developed and characterized properties of new class of binary models where, as observed in biological networks, excitatory neurons are functionally separated from inhibitory units. New patterns, represented as activation and inactivation of binary units in excitatory layer, are stored in the network through recruitment and training of inhibitory units that are group into individual modules and interact with excitatory layer. We investigate the roles the two populations play in memory storage and show that inhibitory layer plays a critical role in memory storage and management and that capacity of this new type of network scales with number of inhibitory neurons. Further, we show that performance of the network is only gradually diminished when excitatory to excitatory connections are removed, but critically depends on inhibitory to excitatory connections. These results are in line with new experimental work showing that inhibitory interneurons are playing critical role in memory storage and recall in the brain networks and may also address why generally excitatory networks exhibit sparser reciprocal connectivity as compared to connections to/from inhibitory units. We further show advantages of so designed coding shame in terms of memory capacity, its expansion with progressive storage of new memories as well as network behavior for large memory loading.

## 1. Introduction

Binary attractor neural networks (ANN) played a critical role in understanding possible dynamical processes underlying standard brain functions such as information storage and retrieval [1; 2; 3]. They are also underlying the Artificial intelligence (AI) revolution happening currently [4; 5; 6; 7]. While number of different variants of these networks were developed, one of the characteristics has been a constant – contrary to known biology, the units in the network have mixed excitatory and inhibitory connections stemming from the same units, forming mixed network layers [2; 8; 9; 10].

Biological neuronal networks stereotypically differentiate into excitatory and inhibitory populations [11], allowing for differential modulation of network dynamics by targeted neuromodulatory inputs [12]. These two populations differ in very fundamental properties. Generally, the excitatory population is significantly larger that the number of inhibitory neurons (with roughly 10:1 ratio) [13]. However interestingly, in biological networks, excitatory neurons are generally sparsely reciprocally connected, while the connections targeting as well as originating from inhibitory interneurons form dense local synaptic arborizations. Subsequently in neocortex and hippocampus excitatory pyramidal cells can sustain firing rate of up to 50Hz but on average fire with frequency below 10Hz [11; 14], whereas inhibitory interneurons encompass many subtypes, but the most numerous groups, parvalbumin positive neurons can support sustained firing frequency of 100Hz or higher [11].

While it has been assumed that both populations are important for memory coding, it was often thought that excitatory neurons carry information content while inhibitory interneurons play more of modulatory role. Recent experimental results have shown however that diverse inhibitory cell populations play critical role in modulation of brain dynamics and function [15]. Interaction of these different classes of neurons is now thought to be critical for memory storage and recall and, further, their relative excitability is thought to play critical role in information processing [15; 16].

Here we construct a binary network model that separates inhibitory and excitatory neurons into two layers, with the memory representations effectively stored withing the patterns of reciprocal inhibitory-excitatory (E-I and I-E) connections. While excitatory layer plays an input/output role, the inhibitory layer has a modular structure which allows for constant expansion and redefinition of the stored memory representations through addition/consolidation of inhibitory modules. We characterize the network properties in terms of storage and retrieval performance in presence of full connectivity and when progressively larger number of connections are missing. We show that the capacity of the network grows directly proportionally to the number of inhibitory cells and while sparsity of excitatory connections only slowly degrades memory performance, the reciprocal inhibitory connectivity is critical for memory storage and recall.

## 2. Results

We develop a binary network framework, where the elements of the network are divided into separate excitatory and inhibitory populations. In this framework excitatory neurons play a role of input pattern detectors as well as network outputs, while the inhibitory layer is divided into modules and the inhibitory units together with their connectivity are critical for pattern storage and separation.

### 2.1. Network dynamics

The network is composed of separate excitatory (E) and inhibitory layer (I), see Fig. 1. The uniform excitatory layer is built of *N*_*e*_ units with its elements being at the same time inputs and outputs of the network. The inhibitory layer is composed of modules which are clusters of inhibitory units associated with the same memory representations. The modules are created as the new binary representations are stored in the network and are composed of one or more inhibitory units.

**Figure 1.**
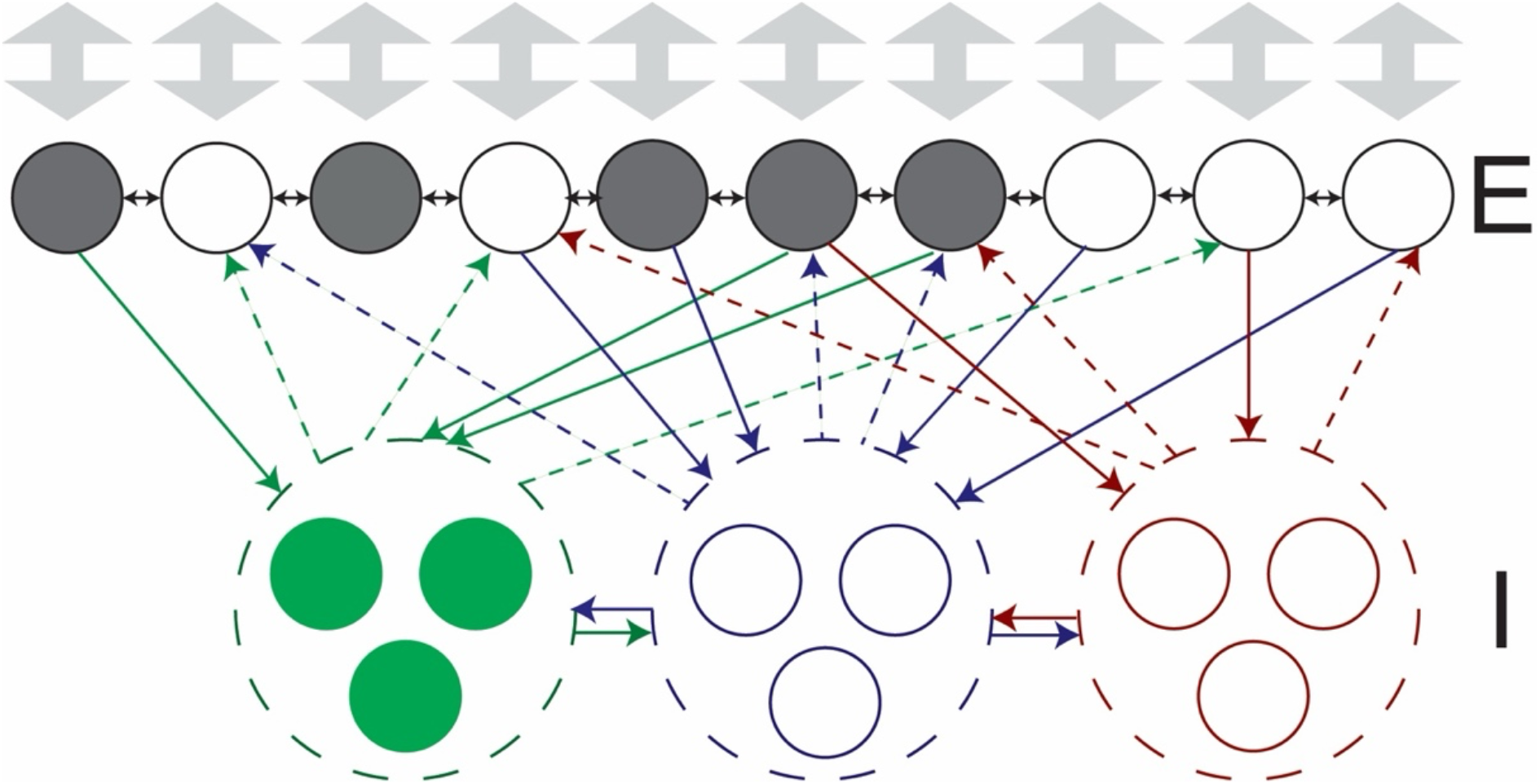
Structure of modular excitatory inhibitory binary network. The excitatory layer is at the same time an input/output layer (gray arrows). The inhibitory modules (dashed circles; here composed of 3 inhibitory neurons each) form as new representations are being stored. Neural representations are composed of *σ*_*j*_ = {0,1} and impinged on the excitatory layer. The excitatory units generate the field, 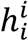, that is activating associated inhibitory units that form modules. These modules on one hand compete with other inhibitory modules for activation (eqn. 4) while on the other control activation patterns of excitatory layer. The inhibitory module with the largest field dominated the dynamics of the network impinging its activation pattern on the excitatory layer. This leads to rapid retrieval of the representation associated with the inhibitory module.

For the excitatory units, following standard approaches [2], we define a signal, 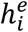, that every excitatory unit receives:

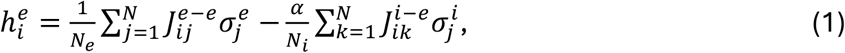

where 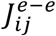 and 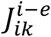 represent input connections to excitatory neurons from other excitatory (referred below as E-E connections) and inhibitory units (i.e., I-E connections), respectively; *N*_*e*_, *N*_*i*_ are numbers of excitatory/inhibitory connections per neuron and *α* in inhibitory connection multiplier.

The excitatory units, similarly to other binary network models, take two states, 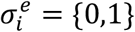, based on the sign local field, 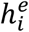 impinging on them:

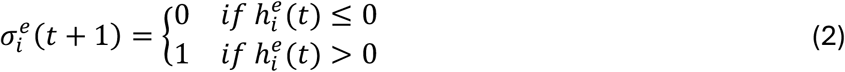

Conversely, the inhibitory layer is composed of modules that receive selective input from the excitatory layer, while their outputs target other modules in inhibitory layer as well as the excitatory layer (Fig. 1). The inhibitory neurons take on values 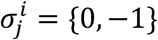.

The inhibitory units within the same modules are not connected but all inhibitory units belonging to different clusters are fully connected creating strong inhibitory interaction between units belonging to different inhibitory modules. Here, the dynamics follows winner-takes-all (WTA) pattern [17; 18] as the neurons within the modules receiving the strongest excitatory field remain active while the others are shut down. Similarly to excitatory units, we define and excitatory field which every inhibitory neuron receives as:

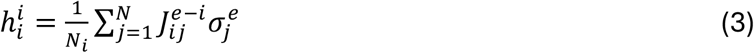

But their state follows the rule:

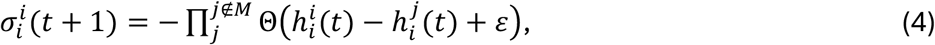

where Θ(·(is a Heavyside function and *ε* is an interaction threshold which allows to control how big the field difference required for the neuron with the lower field to be shut down and can be thought of as an interaction strength between two inhibitory units. Here we set *ε* = 0 for inter-module inhibitory units. At the same time *ε* for intra-module connections is large to ensure that the units belonging to the same module can be coactive. Consequently, the only inhibitory cells that are active at a given time are the ones that have the highest excitatory signal, 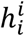, impinging on them. The active inhibitory cells, in turn, inactivate the units in the excitatory layer through E-I connectivity, impinging the pattern associated with the given memory.

#### Network connectivity and memory storage

Memory representations are defined through stable activation patterns of the units in the excitatory layer, 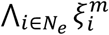, where 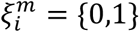 and represents state of i-th excitatory unit in m-th stored representation (memory). Similarly to the classical attractor neural network (ANN) the memory representations are stored in connectivity matrix [2], however the algorithm is adopted to take into account existence and modularity of the inhibitory layer.

The excitatory-to-excitatory (E-E) connections are formed according to standard Hebb’s rule [2], 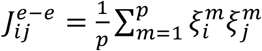. Since 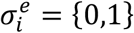, this effectively means that only memory representations having both units active 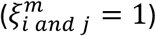 form excitatory connections (see Fig. 1).

The reciprocal connections between the excitatory layer and individual inhibitory modules code the memory representation that is being assigned to the given module. Specifically, all active excitatory units within the given (***m***-th) memory representation, 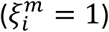, form connections to the inhibitory units composing ***m***-th cluster in inhibitory cell layer (Fig. 1), i.e., 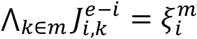, where i denotes i-th excitatory unit and k denotes k-th inhibitory unit belonging to module **m**.

In contrast, inhibitory-to-excitatory connections representing **m**-th memory are formed only from inhibitory units being a part of **m**-th inhibitory module and are targeting excitatory units that are not active in the binary representation assigned to m-th memory (i.e., 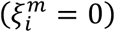, see Fig. 1). The strength of all inhibitory to excitatory (I-E) connections is set to *α* i.e., 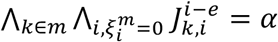. This inhibition is strong, meaning that activated inhibitory unit shuts down targeted excitatory cells, irrespective of the excitatory component of the field that these units receive.

The advantage of such network design is that new representations can be stored progressively in the network without interfering with previously stored memories. During the storage of the new memory representation, the additional inhibitory modules are formed in inhibitory layer as the network is presented with a new memory representation to be stored. Physiologically, these new inhibitory units could be recruited from the pool of unassigned inhibitory cells or alternatively dynamically form new modules from existing but reassigned inhibitory neurons.

New connections are then formed between and within the excitatory and inhibitory layers. Thus, by default the number of excitatory cells remains constant throughout the learning process, whereas the number of inhibitory modules/units grows linearly with the number of stored configurations. The proportionality constant depends on number on units per inhibitory module. We tested performance of the network for modules composed of *S*_*M*_ = 1, 3, 5 inhibitory neurons.

#### Storage capacity

We first investigated the retrieval performance of the network as a function of number of excitatory units, *N*_*e*_. We simulated the network having N_e_=50, 100 and 200 excitatory units (Fig. 2). The inhibitory modules consisted of a single unit and were progressively added with the increasing number of stored binary represetations (i.e., memories). The performance test consisted of retrieval of the memory, with progressively larger fraction (*r = 0*.*1, 0*.*2, 0*.*3, 0*.*4 and 0*.*5*) of units flipped to their opposite state, 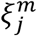 (we will refer to it as representation error). The memory was successfully retrieved when the overlap of the excitatory units in the final network state with given memory 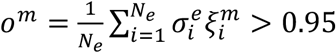. We measure recall fraction which is the fraction of the stored representations recalled correctly, by assigning the value of 1 if the memory was recalled correctly and conversely 0 if it is not. Each point on the Fig. 2 is an average of 20 repetitions.

**Figure 2.**
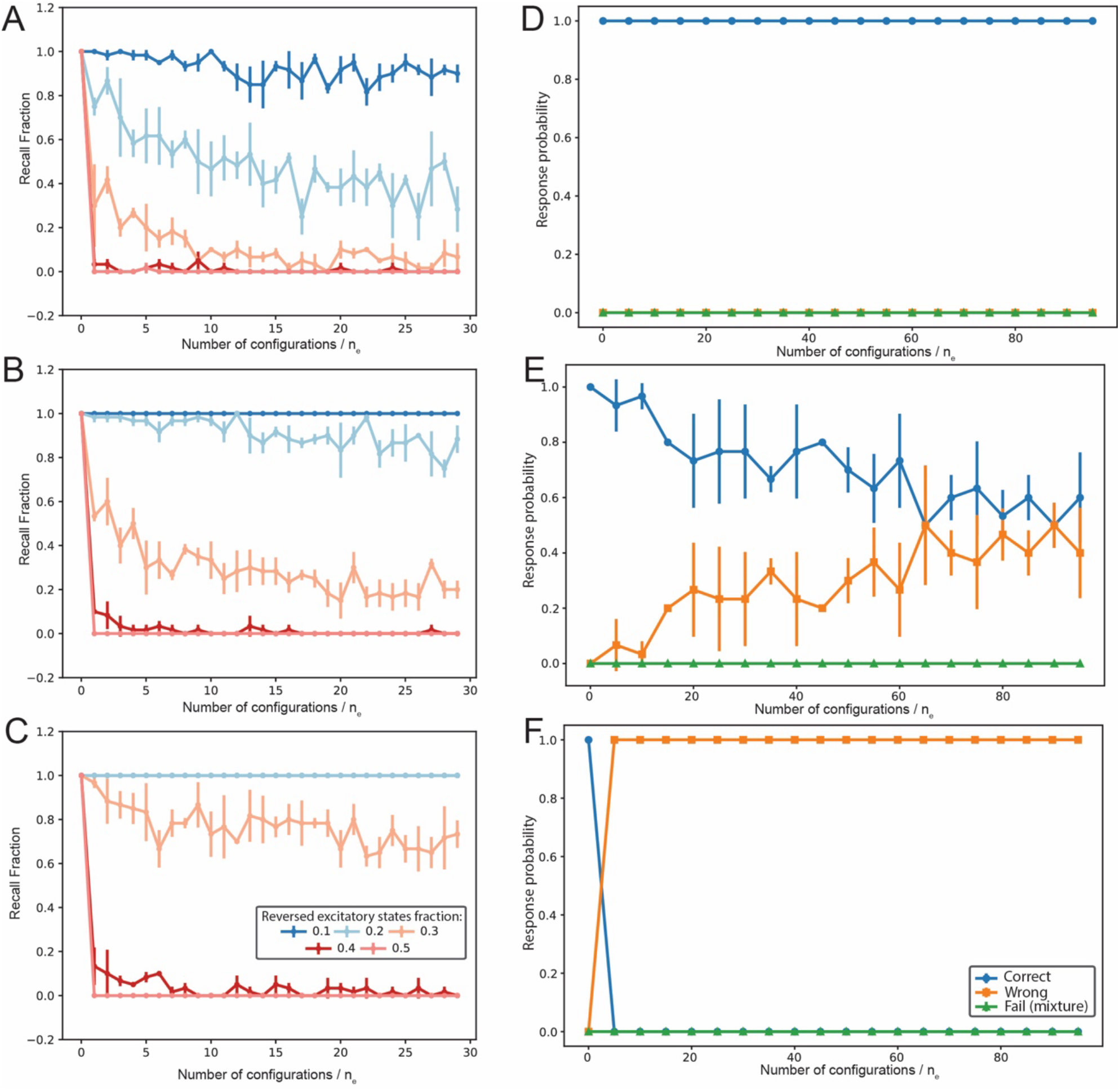
Performance of the network as a function of network size and onset representation error. Performance is measured as a recall fraction (number of correctly recorded configurations/ total number of recalls). Size of the network (number of excitatory neurons): A) N_e_=50, B) N_e_=100, C) N_e_=200. D-F) Type of recall error as a function of the onset representation error for network of N_e_=100. Magnitude of the representation error is: D) 0.1, E) 0.3, F) 0.5. For all simulations the inhibitory modules consisted of single unit. During recall error the network converges toward wrong configuration rather than toward mixed state.

We observe that the network does not saturate and fail even for large number of stored configurations as compared to number of excitatory units (*N*_*e*_). For all network sizes (in terms of the number of units in the excitatory layer) we loaded the networks with up to *p* = 30 · *N*_*e*_ memories, which results in 1500, 3000, and 6000 of configurations stored for n_e_=50, 100 and 200 being, respectively. We have tried larger memory loads and observed stabilization of network performance. We expect however, that the performance of the network will start to deteriorate when the average Hamming distance [2] between the stored representations will get small (and thus overlap between the memories will get large), so that different inhibitory modules will get randomly coactivated due to the small and random field 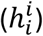 differences, that impinges on inhibitory units.

The performance of the network deteriorates as a function of the representation error (Fig. 2A-C), however it also strongly depends on the number of excitatory units in the network, *N*_*e*_. The representation error of 20% magnitude results in reduction of performance of the network to about 50% for the highest number of memories stored, for the smallest network size with N_e_ = 50 excitatory units. At the same time, 30% representation error reduced network performance only to 70% for network of N_e_=200.

This can be explained by the increasing overlap (decreasing Hamming distance) of the memory representations for the smaller networks and hence by the changes in the mean field magnitude impinged on the inhibitory cells, 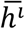, by the excitatory layer. The latter one is critical for activation of correct inhibitory module associated with the given memory (eqn. 4). We therefore investigated how the field depends on, excitatory network size, memory loading (i.e., number of stored configurations) and representation error (Fig. 3). We observed that the mean excitatory field difference between random configuration and the representation stored in the network, does not show strong dependence on memory loading. However, it critically depends on excitatory network size and representation error. Since the selective activation of inhibitory modules, and hence correct memory recall, depends on the separation of the excitatory fields of activated representation from that of all the other stored memories, this separation will play a critical role in memory retrieval.

**Figure 3.**
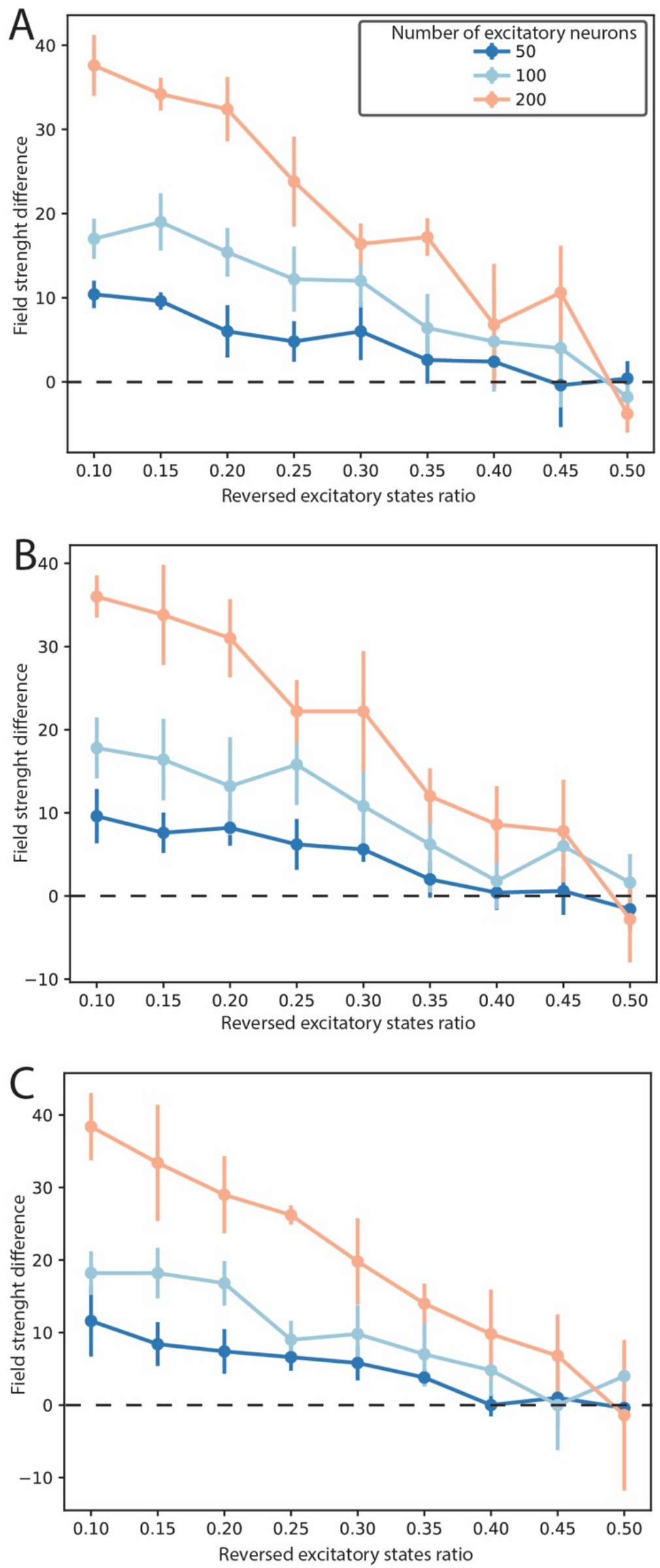
Excitatory field strength difference for various network sizes and as a function of onset representation error (fraction of excitatory neurons with reversed state from stored memory). Ratio of stored configurations to the number of excitatory neurons, 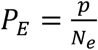, is: A) *PE* = 5, B) *P*_*E*_ = 50, C) *P*_*E*_ = 100. The networks with higher number of excitatory neurons, N_e_, have larger field separation for the same network loading allowing for better memory recall.

We next investigated what types of performance errors dominate during retrieval process. Namely, we were interested in a question that when the network fails to converge to correct memory associated with the given initial state, whether it converged to another stored representation or the system settled in a mixed state with no significant overlap to any of the stored memories. To do so, we measured overlap of the final network state (after the network performed prescribed number of updates) with every stored representation. We divided network responses into three categories: **correct** – when the network converges to the correct memory associated with the initial network state; **wrong** – when the network state converges to another stored representation, and, **fail** - when none of the memories are retrieved (i.e., overlap with all the stored representations stays below recognition threshold). We observed that irrespectively of the applied configuration error (Fig. 3D-F) the network only achieves two states: **correct** recognition or **wrong** convergence. At the same time, as expected, the frequency of **wrong** responses increases with the magnitude representation error.

This result is due to the fact, that the network dynamics is dominated by winner-takes-all effect generated by the inhibitory modules. For high representation errors one of the modules will be selected randomly and the activated inhibitory module will impinge the selected pattern on the excitatory layer leading to recall of one of the stored memories.

#### Network performance upon removal of connections

Next, we characterized network performance when network connections are progressively removed. We removed separately E-E, E-I or I-E connections and measured network recall performance (defined as above) for different number of units in the inhibitory modules. We tested network performance for *S*_*M*_ = 1, 3, 5 inhibitory units in the module.

We observed that the network recall performance was extremely robust to removal of E-E connections (Figs. 4 and 5). Only removal of ∼90% of connections (Fig. 5) made a significant difference in recall statistics.

**Figure 4.**
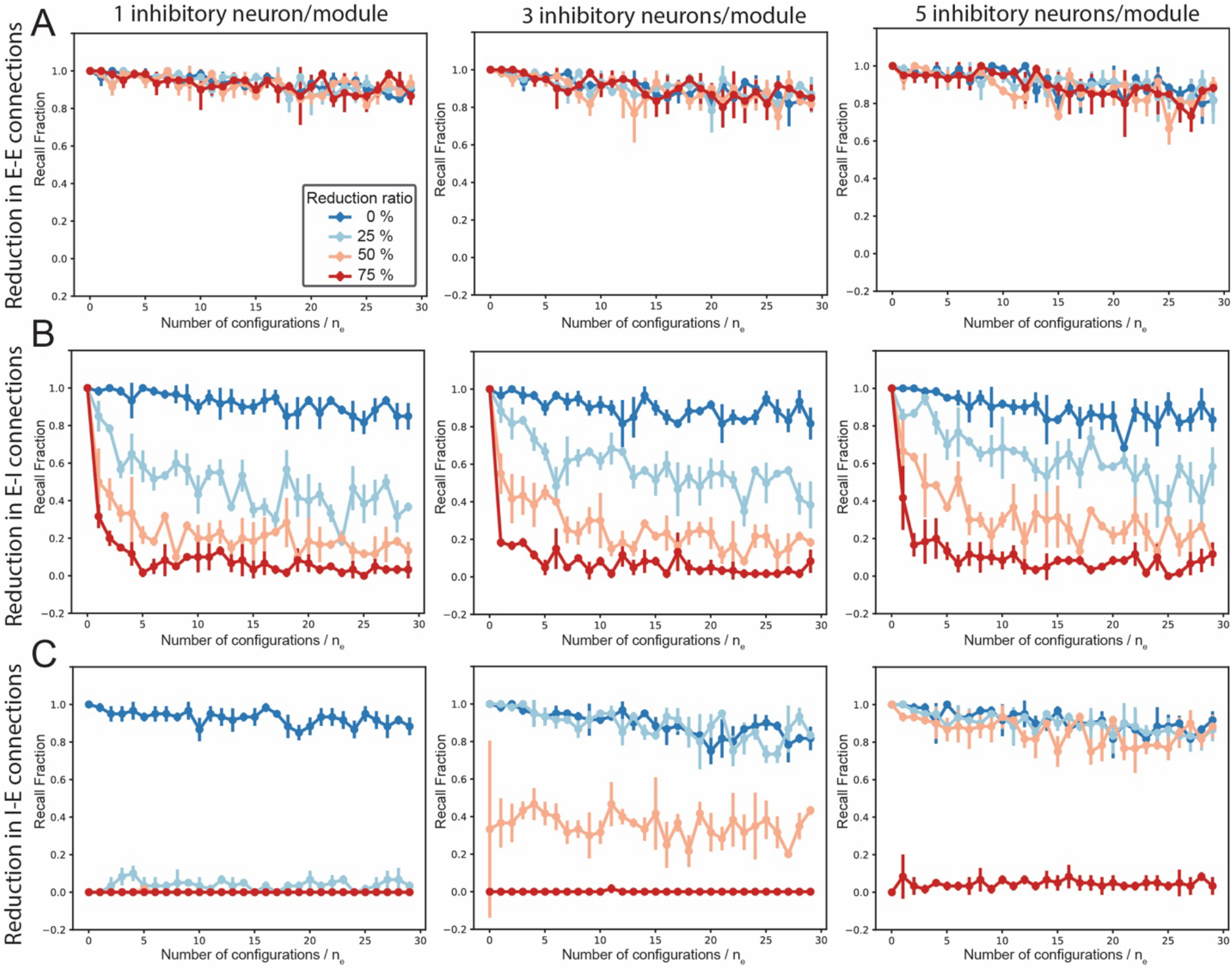
Reduction of network performance (measured as recall fraction) as a function of sparsity of connections for diHerent number of inhibitory units in inhibitory modules. The inhibitory modules consisted of: 1 inhibitory neuron (left column), 3 inhibitory neurons (center column) and 5 inhibitory neurons (right column). In all panels the number of connections was reduced (random removal) by: 0%, i.e., full connectivity (dark blue line), 25% (light blue line), 50% (orange line), 75% (red line). Rows: A) removal of E-E connections, B) E-I connections, C) I-E connections. Network performance is robust to removal of E-E connections independently of number of inhibitory neurons/module, and it is the most sensitive to removal I-E connections. The increase in number of inhibitory neurons per module enhances performance.

**Figure 5.**
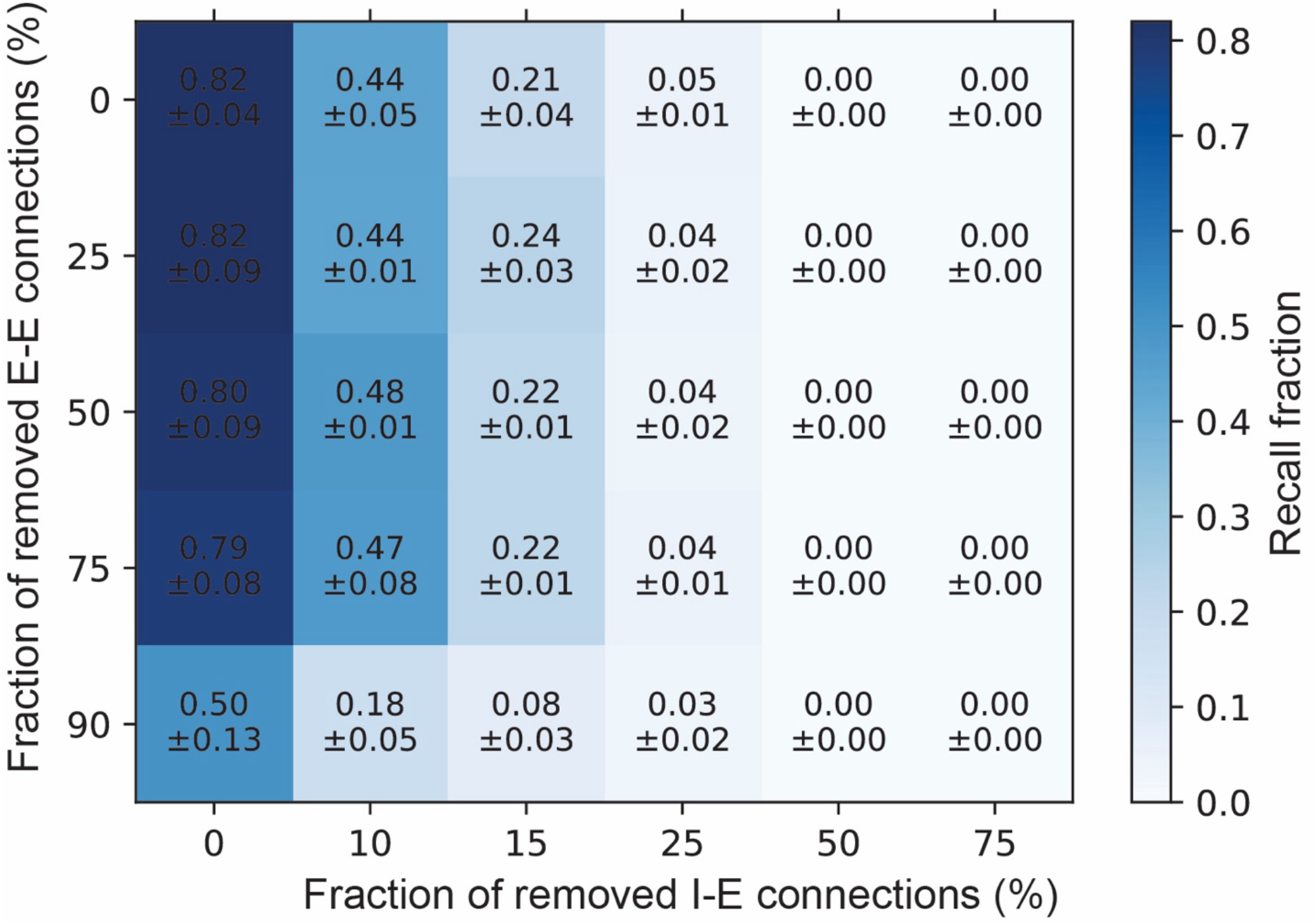
Network performance as a function of fraction of connections removed. Network composed of N_e_=50 with 150 stored configurations is tested, with inhibitory modules consisting of one unit. The results highlight much stronger dependance of network performance on number of I-E connections removed that that when E-E connections are missing.

On the other hand, random removal of E-I connections made a significant difference in network recall as it interferes with the inhibitory neurons’ mean field 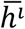, as we observed a precipitous drop in performance for increasing fraction of removed connections. This effect was partially offset by increasing the number of units in inhibitory modules (Fig. 4; center and right columns). The inclusion of five units (*S*_*M*_ = 5(per inhibitory module increased the recall performance by 10-20% for every connection removal fraction (Fig. 4B, right column). This effect is driven by the fact that improved statistics of inhibitory field sampling (due to multiple inhibitory units in a module) could lead to partial recovery of baseline performance. Finaly, the strongest effect was observed in the case of removal of I-E connections (Fig. 4C). Here removal of even low fraction of connections for network composed of inhibitory modules consisting of a single unit resulted in precipitous drop of performance (Fig. 4c, left column). This effect is driven by the fact that I-E connections are solely responsible for impinging the unit inactivation pattern on the excitatory layer. Conversely, missing I-E connections will leave spurious excitatory units activated. Here increasing number of units within the inhibitory module dramatically increases network recall performance as it provides robustness to the targeting of correct excitatory neurons from multiple units within individual inhibitory modules (Fig. 4C; center and right panels).

#### Network performance in the presence of noise

Finally, we investigated how the network preforms in the presence of noise. Following standard approaches [2], the noise is defined as a reduced probability of a unit achieving a state determined by the local field it receives:

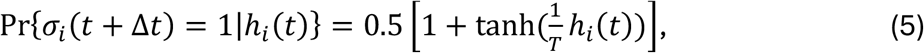

where *T* ∈ [0, +∞) is a noise parameter, with zero denoting, noise free deterministic dynamics.

Here, we added noise only to the excitatory layer, with underlying physiological reason being that excitatory neurons are more prone to noise dynamics because they are directly associated with noisy input and additionally, in the brain their reciprocal (E-E) connectivity is sparse. Inhibitory neurons on the other hand receive more convergent input and form denser arborizations [19] leading to noise cancelation.

We measured how the network recall performance changes as a function memory loading in presence of noise. We varied the noise level between *T* ∈ [0, 0.16]. We observed (Fig. 6), as expected, that network performance declines as the noise level grows, but is largely independent on memory loading (it quickly asymptotes to a constant value for larger memory loading). This is due to independence of inhibitory modules mediating specific memory recall.

**Figure 6.**
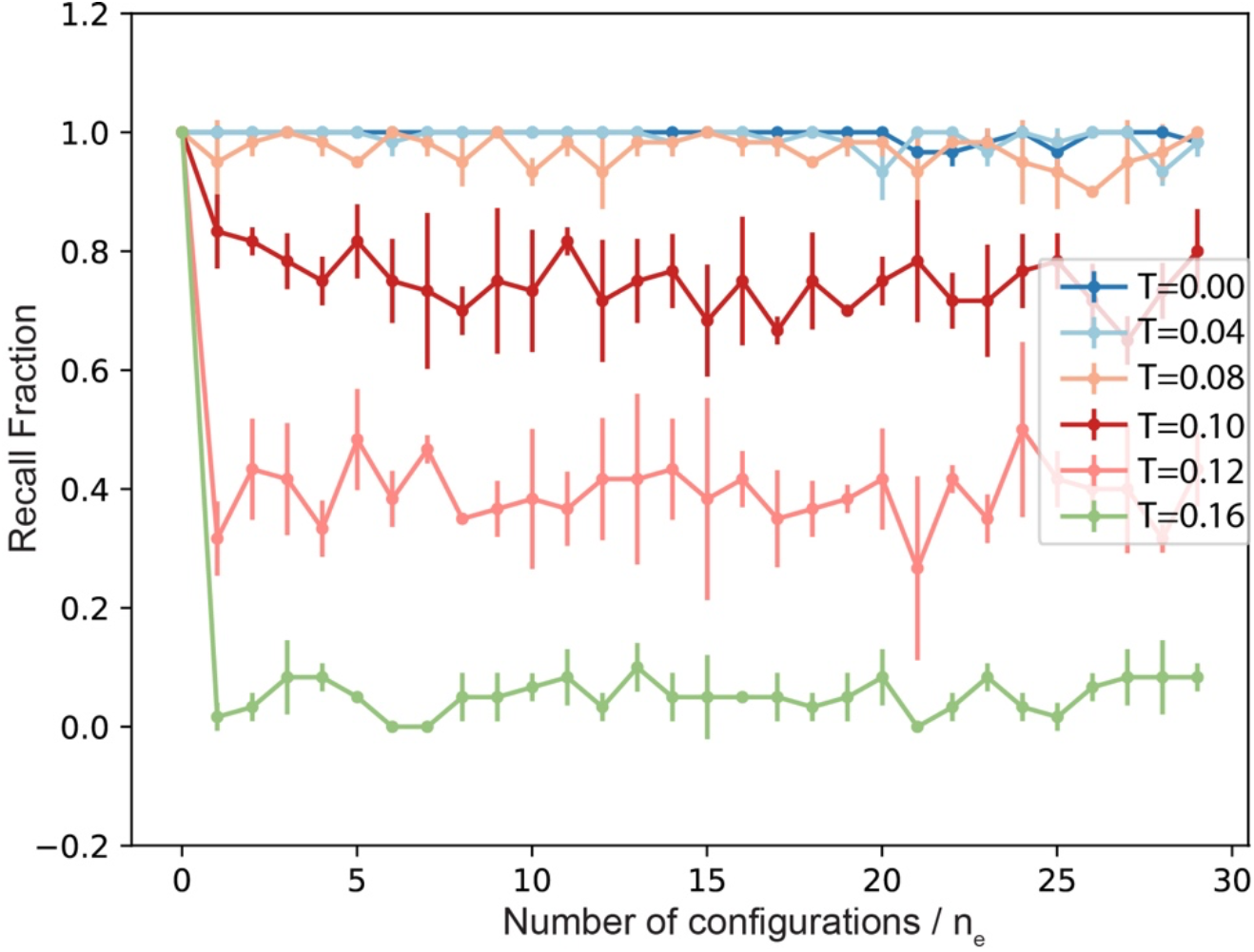
Network performance as a function of noise level (T). The network is composed of N_e_=100 excitatory neurons and with 5 inhibitory neurons per inhibitory module. The network performance in presence of noise does not depend on network loading.

## Discussion and conclusions

In this manuscript we introduce a binary neural network framework that separates excitatory and inhibitory signaling in the network. We argue, that in addition to being more physiologically realistic, the proposed framework has multiple performance advantages to standard, Hopfield type, binary attractor neural networks [1].

In the proposed framework, the excitatory layer carries overall network excitation and plays critical input/output function, while at the same time the information storage and recall is critically controlled by the inhibitory network layer. The inhibitory layer is divided into modules which separately mediate activation/deactivation of the excitatory units which leads to recall of associated memory. The inhibitory units belonging to different modules follow winner-takes-all dynamics allowing for recall of the memory that is most closely associated with the input, while shutting down spurious contribution of other inhibitory modules to the process.

We further show that while the network performance is robust to removal of E-E connections, the E-I connectivity and even more importantly I-E connections play critical role in memory recall.

These properties of closely follow experimental results as well as hypothesized functional properties of the brain networks. It has been shown that the reciprocal (E-E) connectivity between excitatory neurons in the brain is generally very sparse whereas both excitatory to inhibitory (E-I) connections and well as feedback inhibitory-to-excitatory (I-E) connections form dense connectivity trees [19]. At the same time, inhibitory to inhibitory (I-I) connectivity forms dense structures directly on the same interneuron types or indirectly acting through other interneurons [20] [15; 21].

The excitatory representations constituting memories are iteratively stored through recruitment and formation of new independent inhibitory modules. This makes the capacity of the network linearly proportional to number of modules (and thus inhibitory units in the network). Analogously to the results presented here, experimental data analysis and modeling has shown that inhibitory connectivity, rather than excitatory, possibly controls the information maintenance in the cortex [22] and that the number of inhibitory neurons in the network is directly related to maximal number of neural assemblies that can be consolidated, indicating that inhibition has a direct impact on the memory capacity of the neural network [23].

Further, the formation of inhibitory modules can be viewed as emergence of inhibitory engram neurons. Such engrams have been observed in biological networks and, as here, are critical for memory selectivity [17; 24; 25].

Such coding scheme has several additional advantages. We have shown that network can store new configurations progressively as they arrive, independent of its earlier history limiting the interference between the stored representations. Furthermore, because of independence of the module connectivity, the network never experiences catastrophic forgetting [26; 27] associated with overloading.

At the same time, the network can be extended to include progressive forgetting of less frequented memories by progressively weakening excitatory connections to the associated inhibitory modules.

In all, the advantages of the presented binary network framework should to of interest to both neuroscience and AI community.

